# Glia-neuron interactions underlie state transitions to generalized seizures

**DOI:** 10.1101/509521

**Authors:** Carmen Diaz Verdugo, Sverre Myren-Svelstad, Celine Deneubourg, Robbrecht Pelgrims, Akira Muto, Koichi Kawakami, Nathalie Jurisch-Yaksi, Emre Yaksi

## Abstract

Brain activity and connectivity alter drastically during epileptic seizures. Throughout this transition, brain networks shift from a balanced resting state to a hyperactive and hypersynchronous state, spreading across the brain. It is, however, less clear which mechanisms underlie these state transitions. By studying neuronal and glial activity across the zebrafish brain, we observed striking differences between these networks. During the preictal period, neurons displayed a small increase in synchronous activity only locally, while the entire glial network was highly active and strongly synchronized across large distances. We observed that the transition from a preictal state to a generalized seizure leads to an abrupt increase in neuronal activity and connectivity, which is accompanied by a strong functional coupling between glial and neuronal networks. Optogenetic activation of glia induced strong and transient burst of neuronal activity, emphasizing a potential role for glia-neuron connections in the generation of epileptic seizures.

## INTRODUCTION

Epilepsy is a common neurological disorder characterized by recurrent spontaneous seizures (Fisher et al., 2014). When the brain switches from a resting state into a seizure, dramatic transitions occur leading to abnormally excessive or synchronous brain activity. Traditionally, epileptic seizures have been divided into focal and generalized seizures based on their mode of onset. Focal seizures arise from a confined area and may propagate to the rest of the brain (Chang and Lowenstein, 2003; Kramer and Cash, 2012). In contrast, generalized seizures are per definition localized in both brain hemispheres from the onset. However, growing evidence from electrophysiological and functional imaging studies in patients suggests a localized onset also for generalized seizure types (Avoli and Gloor, 1982; Huguenard, 1999; Meeren et al., 2002; Paz and Huguenard, 2015; van Luijtelaar et al., 2014).

Profound changes in neuronal activity and connectivity was observed during state transitions associated with seizure initiation and propagation (van Diessen et al., 2013). High-frequency oscillations and breakdown of inhibition are often considered to be the key triggers of ictal discharges (Bekenstein and Lothman, 1993; Bragin et al., 1999; Cobos et al., 2005; de Lanerolle et al., 1989; Douw et al., 2010; Jiruska et al., 2017; Kramer and Cash, 2012; Paz et al., 2011; Paz and Huguenard, 2015; Prince and Wilder, 1967; Schevon et al., 2012; Trevelyan et al., 2007). Once the inhibition surrounding the epileptic focus breaks down, excessive neuronal activity propagates first locally to highly connected hubs, also called ‘choke points’, and later globally to the rest of the brain (Paz and Huguenard, 2015; Stam, 2016). These hubs display increased synchrony and connectivity during seizures (Douw et al., 2010; Stam, 2016; Yasuda et al., 2015) and are suggested to play an important role in seizure propagation by acting as gatekeepers between seizure foci and the global brain network (Kramer et al., 2008; Morgan and Soltesz, 2008; van Diessen et al., 2013). Interfering with pathological hubs was shown to be effective in controlling seizure propagation (Paz et al., 2013; Sinha et al., 2017; Varotto et al., 2012). Interestingly, focal low amplitude and high frequency neuronal activity evolves into large amplitude and slow frequency activity, as localized seizures propagate (Pinto et al., 2005; Schiff et al., 2000; Schiff et al., 2005). Taken together, the initiation and propagation of seizures is often viewed as transitions of brain states, with prominent changes in neuronal connectivity. For example, epilepsy patients display a transition from a balanced baseline connectivity state into a temporary hypersynchronous connectivity state (Kramer and Cash, 2012; Kramer et al., 2008; Ponten et al., 2007; Schindler et al., 2008). Intriguingly, seizure initiation is also linked to physiological state transitions during sleep (Herman et al., 2001; Hofstra et al., 2009), where changes in neuromodulator levels and neuronal excitability lead to alterations in neuronal connectivity and synchrony (Bazil and Walczak, 1997; Ewell et al., 2015; Malow et al., 2000; Malow et al., 1998; Minecan et al., 2002). It is, however, unclear how such profound changes of neuronal activity and connectivity can occur rapidly during state transitions to generalized seizures.

The important role of glia in modulating synaptic connectivity and neuronal excitability is now well-accepted (Allen and Eroglu, 2017; Bazargani and Attwell, 2016; Perea et al., 2009; Volterra et al., 2014). Astrocytes, the largest group of glial cells, regulate the availability of neurotransmitters and ions at the synaptic cleft, especially glutamate and potassium (Haydon, 2001; Parpura et al., 1994; Volterra and Meldolesi, 2005). Glial signaling is involved in a great variety of brain functions from locomotion to sleep and memory formation (Adamsky et al., 2018; Brancaccio et al., 2017; Nimmerjahn et al., 2009). Moreover, astrocytes are highly connected through gap junctions, and form a functional syncytium, which was shown to effectively redistribute ions and neurotransmitters across large distances in the brain (Scemes and Spray, 2004; Volman et al., 2012). Interestingly, transgenic mice with a deficiency in the astrocytic gap junction coupling show spontaneous seizures (Wallraff et al., 2006). Moreover, alterations of astrocytic coupling and clearance of potassium were linked with temporal lobe epilepsy both in rodents and human patients (Bedner et al., 2015; Schroder et al., 2000; Strohschein et al., 2011; Wallraff et al., 2006). Along with these studies, precisely how glia-neuron interactions are involved in initiation and propagation of epileptic seizures remains an exciting open question with potential therapeutical applications.

To address these questions, we characterized the state transitions of neuronal and glial activity and connectivity in a pentylenetetrazole-induced seizure model of zebrafish larvae (Afrikanova et al., 2013; Baraban et al., 2005; Turrini et al., 2017). The small brains of transparent zebrafish larvae combined with two-photon calcium imaging allowed us to monitor the activity and functional connectivity of thousands of individual neurons and glia, in vivo, with unprecedented spatiotemporal resolution (Wyatt et al., 2015). We showed that during the transition from a baseline state to a preictal state, the neuronal synchrony increased only locally, while the glial synchrony showed an increase across large distances independent of neuronal synchrony and activity. Interestingly, at the initiation of a generalized seizure, when a strong surge of both glial and neuronal activity occurred, we observed a rapid transition in the functional connectivity of neuronal networks and a sudden functional coupling between glia and neurons. Finally, we showed that activation of glial networks leads to a strong increase in neuronal activity. Taken together, these interactions could underlie the rapid transition from a preictal state to a generalized seizure, which goes beyond neuronal connectivity rules. We propose that alterations in glia-neuron interactions can underlie state transitions that lead to generalized epileptic seizures.

## RESULTS

### Calcium imaging is a reliable reporter of neuronal activity related to epileptic seizures

To monitor the seizures across multiple brain regions with high temporal and spatial resolution, we first examined to what extent epileptic activity can be captured with genetically encoded calcium indicators in larval zebrafish. To this end, we measured simultaneously the electrical activity and calcium signals in *Tg(elavl3:GCaMP6s)* (Vladimirov et al., 2014) zebrafish larvae expressing *GCaMP6s* in all neurons (Chen et al., 2013; Reiten et al., 2017). Seizures were induced through application of pentylenetetrazole (PTZ), a widely used proconvulsant agent that is a GABA_A_ receptor antagonist (Afrikanova et al., 2013; André et al., 1998; Baraban et al., 2005; Fisher, 1989; Ono et al., 1990; Schickerová et al., 1984). Previous reports have shown that electrical activity can be reliably measured by local field potential (LFP) recordings at high frequency with a single, intra-encephalic electrode inserted in the midbrain (Afrikanova et al., 2013; Baraban et al., 2005). Hence, we performed LFP recordings in immobilized *Tg(elavl3:GCaMP6s)* larvae under an epifluorescence microscope (Figure 1A). The neuronal activity reported by the calcium indicator clearly reflected the electrical activity measured by LFP (Figure 1B). Importantly, calcium imaging enabled the detection of neuronal events both prior to drug exposure (baseline), prior to the generalized seizure (preictal) and during the generalized seizure, defined as the major epileptic activity spreading to all brain regions simultaneously (Figure 1C). To provide further evidence that calcium imaging can capture changes in the epileptic network activity, we analyzed the oscillatory power of neuronal activity in electrical and calcium signals during these three periods. Since the onset of a generalized seizure varied across larvae, we evaluated the preictal period as the minute preceding the generalized seizure. Power spectral density analysis of LFP and calcium signals showed that the oscillatory changes of network activity can be detected by both techniques and that there was an increase of oscillatory activity during the preictal period at frequencies less than 10 Hz (Figures 1D-1F). During generalized seizures we observed a reduction of power in high frequency activity and increase in power of low frequency activity (Figures 1D-1F). Mean square coherence between LFP and calcium signals indicated that calcium signals can effectively carry information from the preictal and ictal events (Figure 1G). Altogether, our data shows that calcium imaging allows us to detect both the generalized seizures that spread across the entire brain, but also the alterations in neuronal activity during preictal discharges.

**Figure 1.**
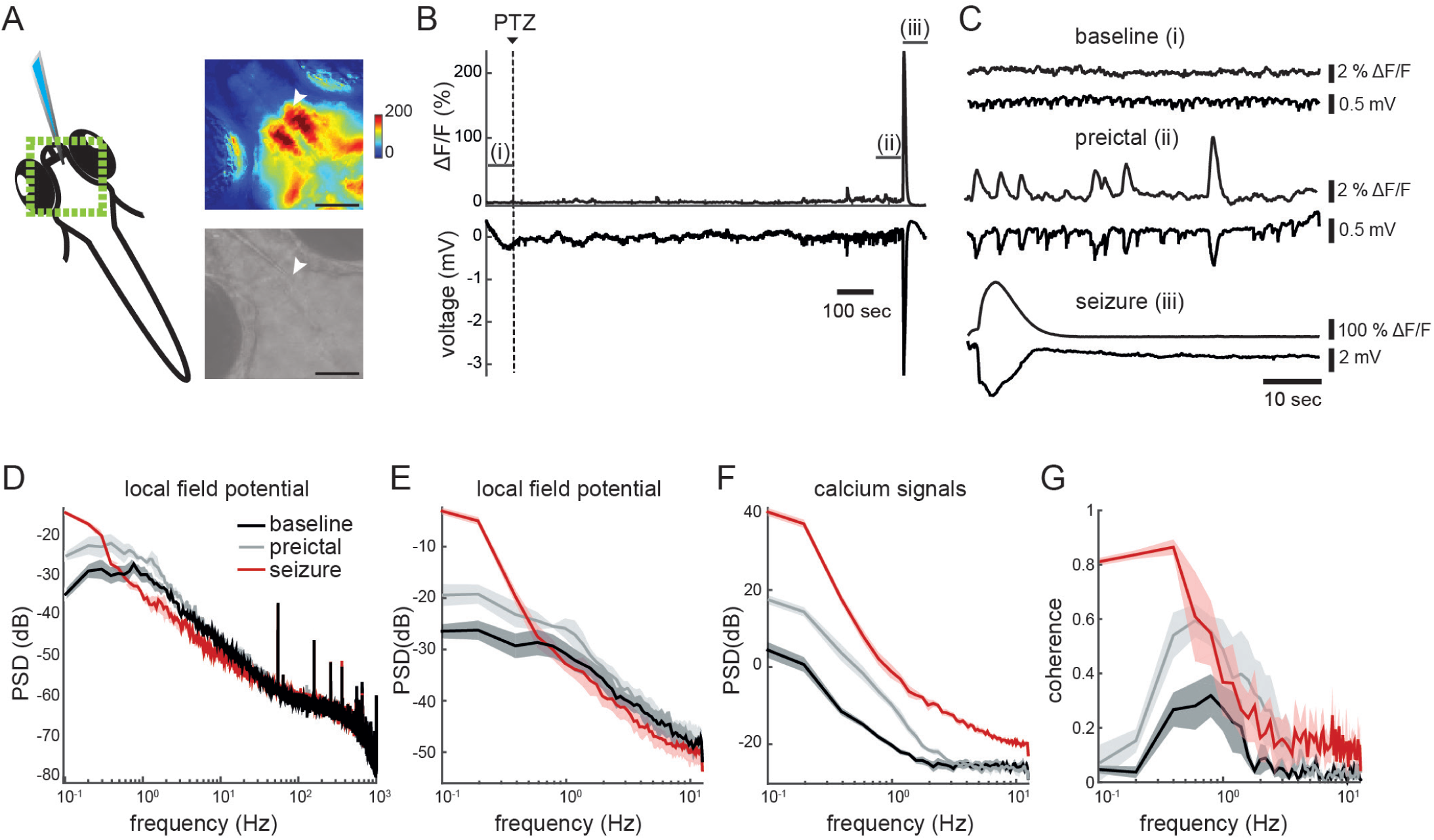
Calcium imaging reliably reports temporal components of neuronal activity during preictal and generalized seizure period. **A**) Scheme representing simultaneous local field potential (LFP) recording and epifluorescence calcium imaging. Green box indicates the region recorded with calcium imaging. The top right image shows the fluorescence intensity change (ΔF/F) during a seizure, warmer colors indicate stronger activity. The bottom gray scale panel is a raw image of a zebrafish brain (7 dpf, *Tg(elavl3:GCaMP6s)*). White arrowhead indicates the electrode position. Black bar reflects 100 micrometers. **B**) Calcium signals (ΔF/F) recorded near the electrode (upper) and electrical activity recorded with the electrode (lower). Dashed line indicates the start of 20 mM PTZ perfusion. **C**) Enlargement of the three time windows marked in Figure 1A. Baseline represents spontaneous activity (prior to drug exposure), preictal period is prior to the onset of the seizure, and the seizure is the first minute following the onset of the generalized seizure. **D**) Power Spectral Density (PSD) of the LFP during the three time windows: baseline (black), preictal (gray) and seizure (red). **E**) PSD of the LFP (decimated to 25 Hz recording rate) **F**) PSD of the ΔF/F calcium signals recorded at 25 Hz. **G**) Magnitude squared coherence between LFP and ΔF/F calcium traces (both at 25 Hz). Shadows represent the standard error of the mean (n=5).

### Brain regions are differentially recruited during epileptic activity

To investigate the neuronal circuits activity during transition of the brain from a healthy baseline state to an epileptic state, we performed two-photon calcium imaging of *Tg(elavl3:GCaMP6s)* zebrafish larvae across multiple brain regions with single cell resolution. We collected data from five anatomically defined brain areas: telencephalon, thalamus, optic tectum, cerebellum and brainstem (Figures 2A-2B, Supplemental Video S1). Upon semi-automated cell detection, we obtained the neuronal activity of more than two thousand neurons in average per fish (2012.6 ± 238.7 (mean ± SEM), n=8) (Figure 2A). To better understand the neuronal network transitions, we first quantified the number of active neurons and their average activity during seizure generation. We observed a small but significant increase in the overall ratio of active neurons during the transition from baseline to preictal period. During the generalized seizure, more than 90% of all measured neurons were active (Figure 2C). Upon analysis of individual brain regions during the preictal period, we saw a significant increase in the ratio of active neurons only in optic tectum and cerebellum, but not in telencephalon, thalamus and brainstem (Figure 2D). Yet, during generalized seizures, all brain regions recruited significantly more active neurons (Figure 2D). Next, we quantified the activity of active neurons by calculating the integral under the curve of the calcium events for individual neurons. While we detected an activity increase of more than tenfold in all brain regions during generalized seizures (Figure 2E), throughout the preictal period we observed a significant increase in the activity of active neurons only in the optic tectum (Figure 2F). Taken together, our results showed that individual brain regions are differentially recruited during the preictal period, with significantly larger changes in the optic tectum and the smallest changes in the telencephalon. This is in line with the higher ratio of GABAergic cells (*Tg(gad1:GFP*)) (Satou et al., 2013) vs glutamatergic cells (*Tg(vglut2a:dsRED*)) (Miyasaka et al., 2009) that we observed in the optic tectum, when compared to telencephalon (Figure 2G). Our results indicated that the relative amount of inhibitory cells relates to differential recruitment of distinct brain regions during epileptic activity, and interfering with inhibition in the GABA rich brain regions can lead to triggering of preictal processes.

**Figure 2.**
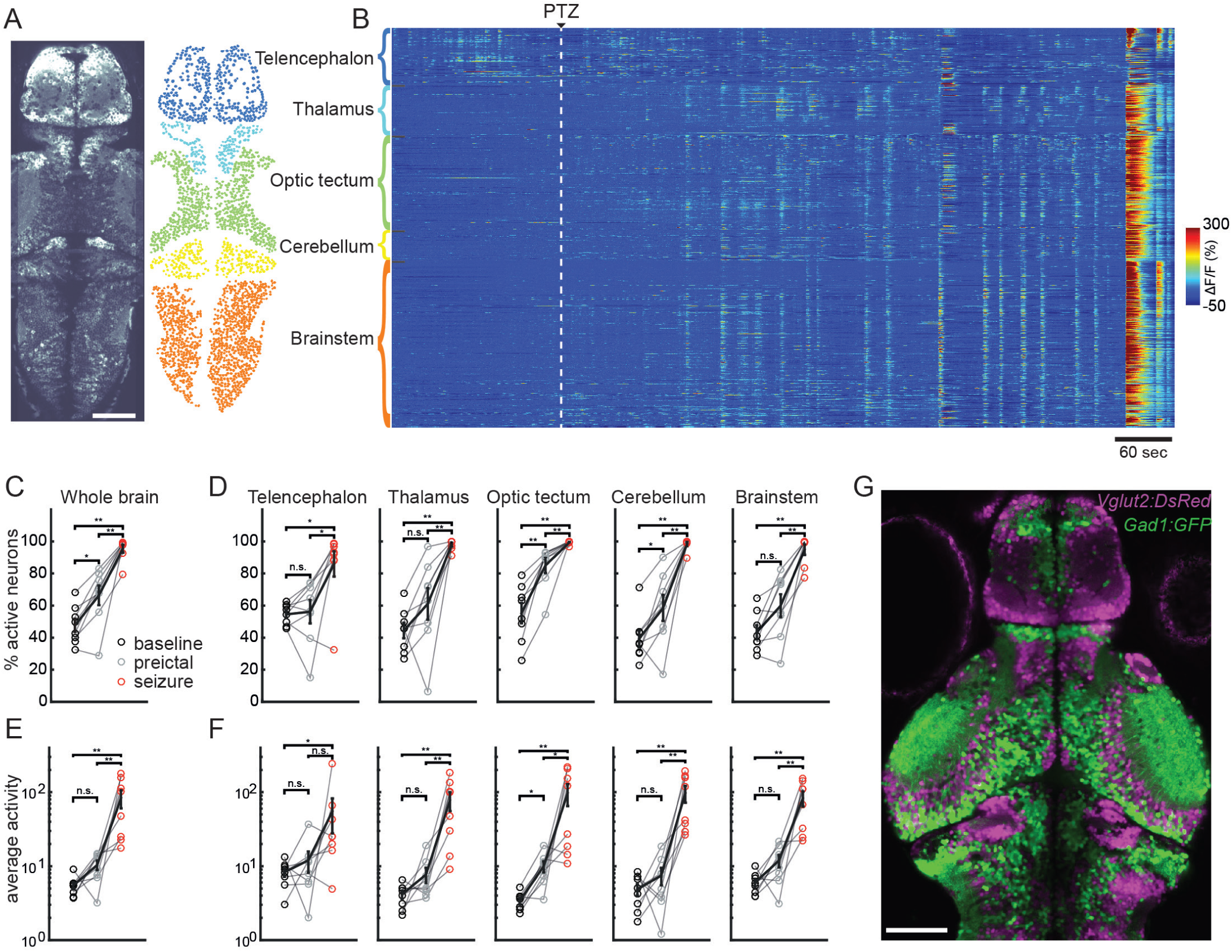
Brain regions are differentially recruited during epileptic activity. **A**) An optical section of a zebrafish larva expressing GCaMP6s in all neurons, obtained by two-photon microscopy, dorsal view (left). Individual neurons (right) in color-coded brain regions: telencephalon (dark blue), thalamus (light blue), optic tectum (green), cerebellum (yellow), and brainstem (orange). **B**) Activity of individual neurons (ΔF/F) over time, organized by brain region. White dashed line indicates the start of 20 mM PTZ perfusion. Warmer colors indicate stronger activity **C**) Percentage of active neurons (>3std_baseline_) in the brain. **D**) Percentage of active neurons per brain area. **E-F**) Average activity of the active neurons, defined by the area under the curve of the ΔF/F trace, in the whole brain (E) and per brain area (F). n = 8 fish (***p= <0.0005, **p= <0.005, *p= <0.05, ns= not significant, Wilcoxon signed-rank test). **G**) Confocal image of *Tg(gad1:GFP)* and *Tg(vglut2a:dsRED)* double transgenic zebrafish larva showing glutamatergic (magenta) and GABAergic (green) neurons. White bar reflects 100 micrometers.

### The abrupt transition to a generalized seizure overrules the functional connectivity

The current knowledge suggests that a switch from a balanced connectivity state into a hypersynchronous connectivity state underlie generation of epileptic seizures. To test this hypothesis, we investigated the alterations in functional connectivity within and across brain regions during preictal and ictal periods. To quantify the functional connectivity, we calculated the pairwise Pearson’s correlations between neurons throughout seizure generation. We showed that during generalized seizures, as the entire brain became active, neurons were highly correlated to each other across brain regions (Figures 3A-3B). But surprisingly, we did not observe a significant increase in correlation between baseline and preictal period while comparing all neurons across the brain (Figure 3B). When brain regions were analyzed individually, there was a small but significant increase in correlations between neurons located within the optic tectum, cerebellum and brainstem (Figure 3B). Across brain regions we showed increase in correlations between thalamus and cerebellum, as well as between optic tectum and brainstem (Figure 3C). During a generalized seizure all neurons and all brain regions were synchronized (Figures 3B-3C). These findings are in line with the hypothesis that nearby neurons would be more likely to have functional connections, thereby showing stronger synchrony during baseline and preictal period. To directly test the relationship of distances between neurons and their functional connections, we visualized the correlations of neuronal activity with neuronal distance. Already during the baseline period, nearby neurons showed stronger correlations (Figure 3D, black solid lines) when compared to spatially shuffled distributions, where neuronal positions were randomly assigned (Figure 3D, black dotted lines). These results confirm stronger functional connectivity between nearby neurons and weaker connectivity between neurons that are more than few hundred micrometers away. During the preictal period, the synchrony between nearby neurons increased, in line with increased connectivity and synchrony within several individual brain regions shown in Figure 3B. On the contrary, the synchrony between neurons that are more than only few hundred micrometers apart showed no or little change of connectivity and it was close to the connectivity levels of spatially shuffled neurons (Figure 3D, gray lines). During the generalized seizures, however, we observed a dramatic increase in neuronal synchrony even between those neurons that are more than several hundred micrometers apart (Figure 3D, red lines). These findings suggest a drastic alteration of neuronal connectivity and synchrony during generalized seizures that is largely independent of spatial distances between neurons, which might be mediated by processes beyond typical neuron-to-neuron communication.

**Figure 3.**
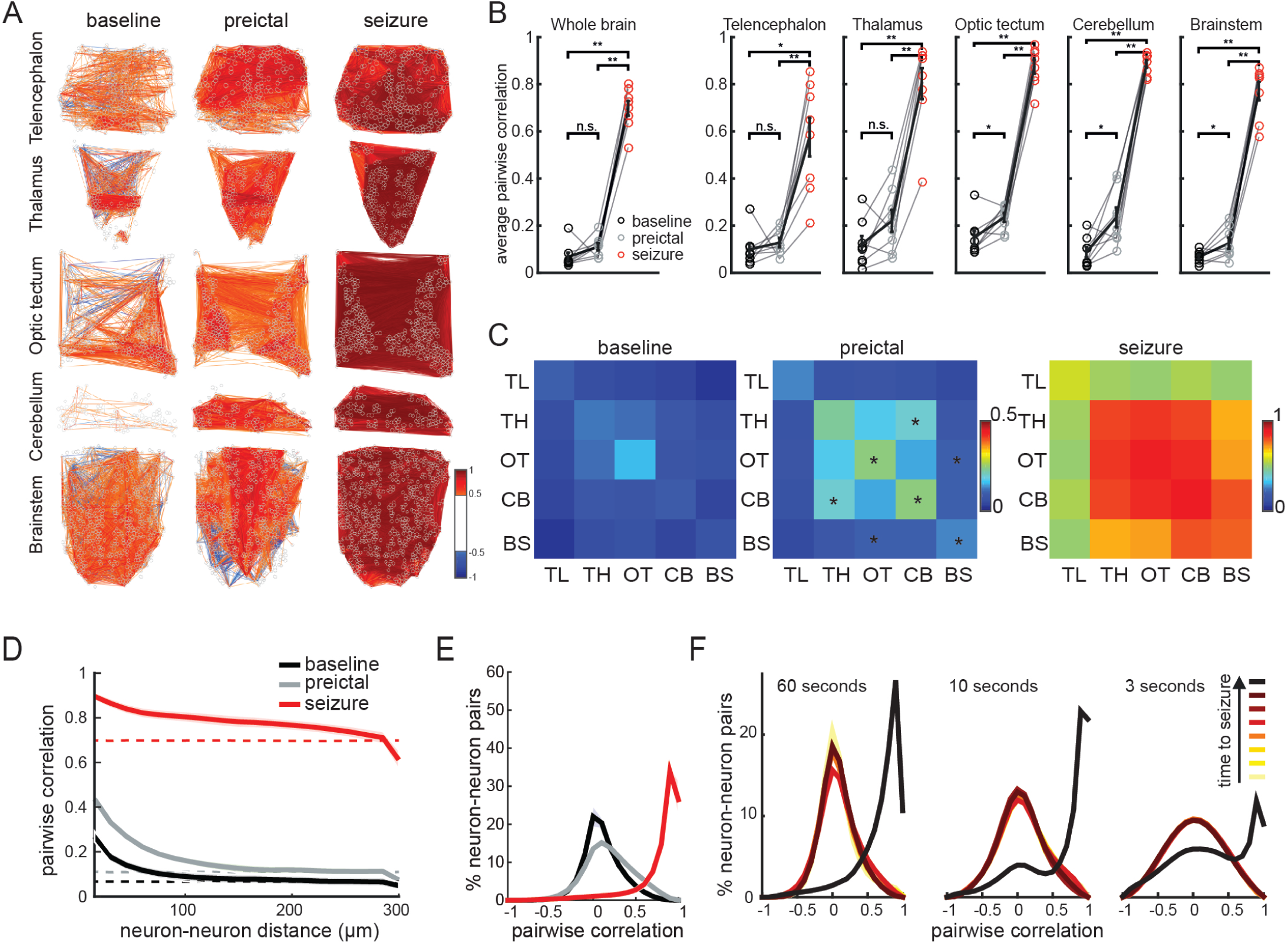
Functional connectivity between neurons changes abruptly from the preictal period to the generalized seizure. **A**) A map depicting pairwise Pearson’s correlation coefficients of neuronal activity within brain regions during baseline, preictal and seizure periods. Each colored line indicates a strong positively (>0.5 in red) or negatively (<-0.5 in blue) correlated activity between pairs of neurons located at the end of the lines. **B**) Average pairwise Pearson’s correlation during baseline (black), preictal (gray) and seizure (red) periods across the whole brain, and within individual brain regions. **C**) Correlation matrices indicating average pairwise Pearson’s correlations of neuronal activity across brain regions, during baseline, preictal and seizure. (Telencephalon, TL; thalamus, TH; optic tectum, OT; cerebellum, CB; and brainstem, BS). Warm colors indicate stronger positive correlations. **D**) Relation between pairwise correlation of neuronal activity and the distance between each neuron pair. Dotted lines represent the results when neuronal locations are shuffled. **E**) Histogram representing the distribution of all correlation coefficient between neurons from all animals during baseline (black), preictal (gray) and seizure periods (red). **F**) Correlation coefficients during seven time periods immediately preceding a generalized seizure. The time periods are of 60 sec, 10 sec, and 3 sec length, respectively. Darkening of colors indicates temporal proximity to the seizure. n = 8 fish (***p= <0.0005, **p= <0.005, *p= <0.05, ns= not significant, Wilcoxon signed-rank test).

To further investigate this drastic alteration of neuron-to-neuron communication, we studied the nature of functional connectivity between all neurons across the brain by visualizing positive and negative correlations in the form of a histogram. In line with our findings based on average correlations across all neurons (Figure 3B), we observed only a slight increase in positive pairwise correlations between neurons from the baseline to the preictal period (Figure 3E). To our surprise, during generalized seizures most negative correlations between neurons were eliminated, indicating a major rearrangement of neuronal connectivity rules. We hypothesize that such a rearrangement of neuronal connectivity could in principle be either due to gradual elimination of inhibitory connections through the antagonistic action of PTZ on GABA_A_ receptors or due to processes that inject massive excitation into the neuronal network and synchronize even those neurons with inhibitory connections by simultaneously activating them. To investigate the time scale for this drastic alteration of neuronal connectivity, we measured the neuronal synchrony at different time points (seven intervals of 60 sec, 10 sec or 3 sec) before the seizure. We found that the drastic rearrangement of neuronal synchrony was not a gradual process that is progressively morphing the neuronal connectivity rules, but instead a rapid process that reshapes the functional connectivity histograms, within less then few seconds (Figure 3F). These results suggest that it is highly unlikely that the alteration of neuronal connectivity is due to gradual elimination of inhibitory connections. Instead, our data was consistent with a rapid process that can deliver excitation to the entire nervous system independent of spatial constraints associated with neuronal connectivity rules.

### Glial cells are highly active and strongly synchronized during preictal period

The rapid and non-spatially constrained transition of neuronal activity and connectivity during seizure generation suggests that the underlying mechanisms are beyond neuronal connectivity rules mediated by classical synaptic transmission. These findings further highlight a potential role for non-neuronal cells in the brain, such as glia, for the spreading of seizures. To study glial activity during seizure development, we generated a *Tg(GFAP:Gal4)nw7* zebrafish line expressing *Gal4* under the astrocytic *glial fibrillary acidic protein (GFAP)* promoter (Brenner et al., 1994; Shimizu et al., 2015). In zebrafish, *GFAP* promoter was shown to label radial glial cells, which in addition to their neurogenic capacity (Hansen et al., 2010; Shimizu et al., 2015; Silbereis et al., 2016), serve the function of mammalian astrocytes (Grupp et al., 2010; Lyons and Talbot; Papadopoulos and Verkman, 2013). We first demonstrated that *GFAP:Gal4* expressing cells are primarily glia and not neurons. To this end, we crossed *Tg(GFAP:Gal4)nw7;Tg(UAS:GCaMP6s)* fish with *Tg(elavl3:jRCaMP1a)* (Dunn et al., 2016) fish expressing the red-shifted calcium indicator *jRCaMP1a* in all neurons. Confocal images revealed exclusively *GFAP* expressing cells along the ventricular zones at the midline, whereas we observed more overlap with *elavl3* positive neurons in other parenchymal zones (Figure 4A). To solely study the activity of *GFAP* expressing glial cells, we focused our analysis only to the region along the ventricles with no neuronal labelling. Next, we performed two-photon calcium imaging on *Tg(GFAP:Gal4) nw7;Tg(UAS:GCaMP6s)* zebrafish larvae, where PTZ was added after a baseline period, as described earlier with *Tg(elavl3:GCaMP6s)* fish. We observed strong increase in glial calcium signals after the perfusion of PTZ (Figure 4B and Supplemental Video S2) that was clearly distinct from the neuronal activity we recorded (Figure 2B). We noticed large, long lasting and highly synchronous glial calcium events in the preictal period, as well as very large calcium increase during generalized seizure (Figure 4B and Supplemental Video S2). Upon quantification of glial calcium signals, we measured significant increases in the percentage of active glial cells (Figure 4C) and in the average activity of glial cells (Figure 4D). During preictal period, in contrary to neurons, glial cells displayed a large and significant increase in synchrony (Figure 4E) that spread across large distances (Figure 4F). During generalized seizures, all glial cells were highly synchronized (Figure 4E) across the brain (Figure 4F), similar to neurons (Figure 3D). Finally, we observed a more gradual increase of glial synchrony during preictal period (Figures 4F and 4G), which suggest an alteration of activity and connectivity of glial networks preceding generalized seizures. Taken together, our results showed that glial cells are highly interconnected and recruited by the nervous system during the preictal period, highlighting an important difference between the glial and neuronal activity preceding generalized seizures.

**Figure 4.**
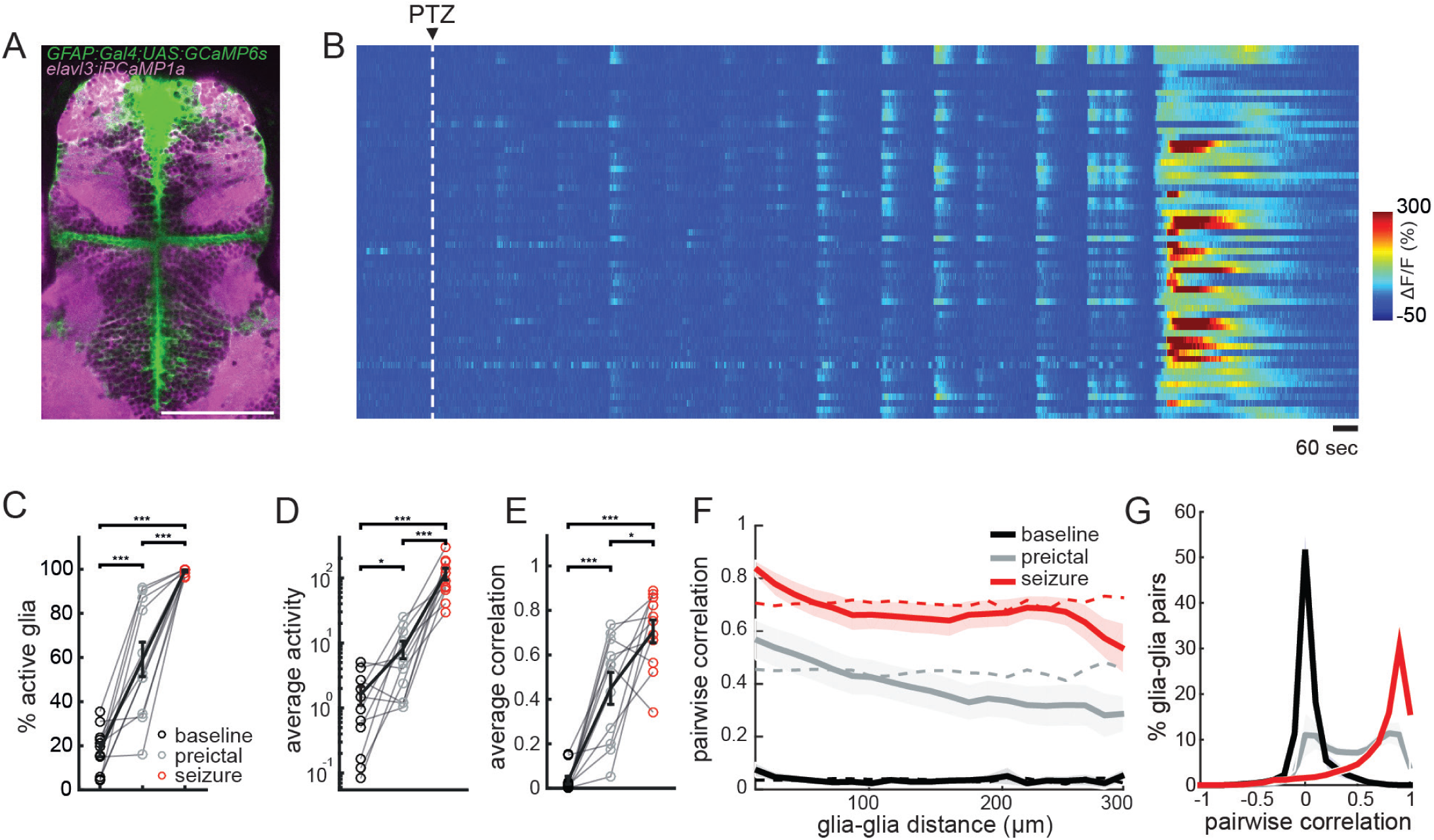
Glial cells are highly active and strongly synchronized already during preictal period. **A**) Confocal image of a transgenic zebrafish larvae expressing *GCaMP6s* in *GFAP* positive glial cells (green) and jRCaMP1a in *elavl3* positive neurons (magenta), dorsal view. White bar reflects 100 micrometers. **B**) Glial calcium signals measured by two-photon microscopy in *Tg(GFAP:Gal4)nw7;Tg(UAS:GCaMP6s)* transgenic zebrafish larva. White dashed line indicates the start of 20 mM PTZ perfusion. **C**) Percentage of active glial cells (>3std_baseline_) during baseline (black), preictal (gray) and seizure (red) periods. **D**) Average activity of the active glial cells, defined by the area under the curve of the ΔF/F trace. **E**) Average pairwise Pearson’s correlation between glial cells **F**) Relation between pairwise correlation of glial activity and the distance between each glial pair. Dotted lines represent the results when glial locations are shuffled. **G**) Histogram representing the distribution of the correlation coefficients between glial cells during baseline (black), preictal (gray) and seizure periods (red). n = 11 fish (***p= <0.0005, **p= <0.005, *p= <0.05, ns= not significant Wilcoxon signed-rank test).

### Glia-neuron interactions change drastically during generalized seizures

Our analyses of glial activity suggest an important role for glia during the transition of preictal activity to a generalized seizure. To investigate the functional interactions between glia and neurons, we simultaneously measured glial and neuronal calcium signals. To achieve simultaneous labelling of glia and neuron, we injected *Et(−0.6hsp70l:Gal4-VP16) s1020t;Tg(UAS:GCaMP6s)* zebrafish expressing *GCaMP6s* in only thalamic neurons with *GFAP:Gal4* plasmid at the one-cell stage. This approach allowed us to visualize the calcium signals of both thalamic neurons and sparsely labelled *GFAP* expressing glial cells along the ventricles (Figure 5A). Consistent with our previous data (Figures 3-4), we observed a strong and synchronous glial activity before and during generalized seizure (Figures 5B, 5C and 5D), while the neuronal activity and synchrony was strongly increased only during generalized seizure (Figures 5B, 5C, and 5D). To investigate functional interactions between glia and neurons we calculated pairwise Pearson’s correlations between every glia and neuron. Interestingly, we observed that the synchronization of glial and neuronal activity drastically increased only during generalized seizures (Figure 5D). These results highlight that during the preictal period, glial activity and synchrony increases drastically but with little functional coupling with neuronal networks. Whereas during generalized seizures, we observed a very strong functional coupling between glial and neuronal networks, emphasizing a potential role for glia-neuron connections in the spreading of ictal activity across large distances in the brain.

**Figure 5.**
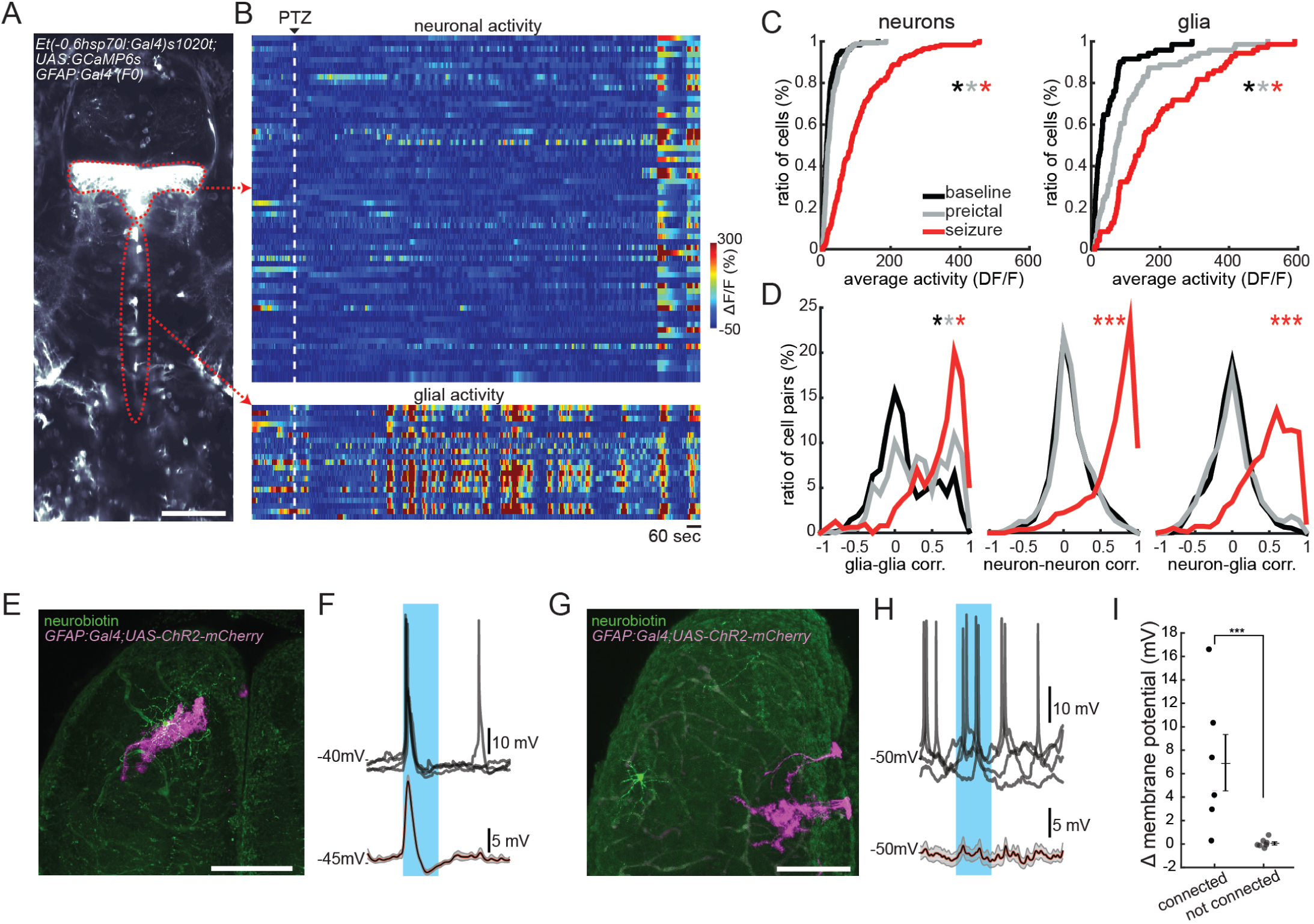
Functional interactions between glia and neurons change drastically during seizure generation. **A**) An optical section of a transgenic zebrafish larva expressing *GCaMP6s* in thalamic neurons (top red dotted line) and *GFAP* expressing glial cells near the ventricles (bottom red dotted line) obtained by two-photon microscopy, dorsal view. White bar reflects 100 micrometers. **B**) Activity (ΔF/F) of individual thalamic neurons (top) and glial cells along the ventricle (bottom). White dashed line indicates the start of 20 mM PTZ perfusion. Warmer colors indicate stronger activity. **C**) Cumulative distribution of neuronal (left) and glial activity (right) during baseline (black), preictal (gray) and seizure (red). **D**) Histograms representing the distribution of all pairwise Pearson’s correlations for the activity of glia-glia pairs (left), neuron-neuron pairs (middle) and glia-neuron pairs (right). **E) and G)** Dorsal view of brain-explant preparation from zebrafish that are sparsely expressing channelrhodopsin2 in *GFAP* positive glial cells (magenta), and randomly recorded individual neurons that are filled with neurobiotin (green) during intracellular recordings. **F**) Membrane potential of an example neuron that has processes passing nearby a patch of channelrhodopsin2 expressing glial cells in (E), during 1 second blue light stimulation (blue shade). Note the strong transient activation of the neuron, in overlaid individual sweeps (top) or 30 Hz low-pass filtered and averaged membrane potential (bottom). **H)** Membrane potential of another example neuron that has processes far away from a patch of channelrhodopsin2 expressing glial cells in (G). Note that the optogenetic activation of glial cells has no effect on the spiking (top) or average membrane potential of this neuron (bottom). **I)** Average change in the membrane potential of neurons during the first 300 milliseconds of optogenetic stimulation of glial cells. Note that the neurons with arborizations that are physically connected with channelrhodopsin2 expressing glial patches display large increase in membrane potential. Whereas neurons with no connections to glial arborization showed no change in membrane potential. n=13 neurons from 13 fish (***p= <0.0005, Wilcoxon ranksum test).

One potential mechanism for synchronized glial activity to trigger a generalized seizure might be through a direct excitation of neuronal networks. Such an excitation by glia onto neurons could in principle be delivered across large distances through gap junction coupled glial networks, which can explain why neuronal networks do not follow functional connectivity rules during generalized seizures. To test whether the activation of glial cells can indeed trigger significant excitation of nearby neurons, we recorded intracellular membrane potential of neurons, while optogenetically activating glial cells sparsely expressing channelrhodopsin-2 in *Tg(GFAP:Gal4);Tg(UAS:ChR2-mCherry)* zebrafish line (Figure 5E). These recordings showed that activating only few glial cells can generate a transient but strong excitation of nearby neurons (Figures 5F-5I). Interestingly, we also observed that glia-to-neuron connections are present only if neurons have processes nearby glial cells that are anatomically confirmed by dye fills. Those neurons that do not have processes nearby sparsely labelled glial patches showed no significant change in their membrane potential (Figures 5G-5I). Taken together, our results revealed that glial activation can strongly modulate the activity of neurons by delivering transient but strong excitation. Such modulation could underlie the strong coupling between glial and neuronal networks during generalized seizures (Figure 5D).

## DISCUSSION

Our results revealed that the transition from a preictal state to a generalized seizure state is an abrupt process accompanied by a strong alteration of the functional connectivity between glial and neuronal networks of the brain. Thanks to the small size of the zebrafish brain, we could follow the seizure development in thousands of individual neurons and glia across multiple brain regions. Such a combination of scale and detail across many brain areas would be difficult in studies of epilepsy in mammals. Using simultaneous electrical recordings, we confirmed that genetically encoded calcium indicators in combination with two-photon imaging can reliably report epileptic activity, especially on the lower frequency ranges related to preictal and ictal events. Hence, we propose that two-photon calcium imaging is complementary to commonly used high-throughput behavioral assays in zebrafish for studying mechanism underlying seizure generation (Afrikanova et al., 2013; Baraban et al., 2005; Samarut et al., 2018).

We provide evidence for several common physiological features between zebrafish and mammalian models of epilepsy. Previous studies suggest that the seizure propagation arises from profound alterations in local and global network connectivity (Kramer et al., 2008; Morgan and Soltesz, 2008; van Diessen et al., 2013). In our PTZ induced seizure model, we observed recruitment of more neurons as the network shifts from baseline to preictal state. This was also accompanied by a small and brain-region-specific increase of neuronal synchrony between nearby neurons. Consistent with previous reports (Dichter and Spencer, 1969; Paz and Huguenard, 2015; Prince and Wilder, 1967; Trevelyan et al., 2007), we observed that this enhanced preictal neuronal activity and synchrony was mainly confined to inhibitory-neuron rich brain regions, where inhibitory surround broke down upon exposure to the GABA_A_ antagonistic proconvulsant PTZ. Interestingly, among the five major brain regions that we monitored, the optic tectum and the cerebellum were the most prominent brain regions to show significant alterations during the preictal period, followed by the thalamus and the brainstem. The telencephalon instead was recruited only during generalized seizures. This sequential recruitment of brain areas could relate to different ratio of excitatory and inhibitory neurons. Indeed, our results suggest that the optic tectum, which is more susceptible to PTZ treatment, has larger ratio of GABAergic neurons than the telencephalon with lowest ratio of GABAergic vs glutamatergic neurons, similar to mammalian cortex (Mueller et al., 2006). Alternatively, it was proposed that the spreading of epileptic activity may require involvement of epileptogenic hubs, also known as choke points (Paz and Huguenard, 2015; Stam, 2016). As such, these confined areas, which have profound connections to the rest of the global network would allow oscillatory activity to spread from local to global networks, resulting in generalized seizures. Similar to the mammalian brain, our results suggest that the zebrafish midbrain regions such as the thalamus and the optic tectum (homologous to mammalian superior colliculus) appears to have hub-like properties with earlier recruitment of synchronous activity before generalized seizures. These findings highlight a potential role of non-cortical brain regions in the initiation and spreading of seizures. Interestingly, the first epilepsy with a proven monogenic etiology, autosomal dominant sleep-related hypermotor epilepsy (ADSHE), is accompanied by prominent alterations in these midbrain regions in human patients (Picard et al., 2006). Future studies on the role of non-cortical brain regions in human epilepsies will likely provide a better understanding of underlying mechanisms.

Our results revealed that the abrupt transition from a preictal state to a generalized seizure was accompanied by a drastic reorganization of functional connectivity and synchrony across neurons. Similar to epilepsy patients and rodent models (Pinto et al., 2005; Schiff et al., 2000; Schiff et al., 2005), our electrical LFP recordings in zebrafish brain suggest a reduction of high frequency activity and an increase in low frequency activity during seizure generation. We observed that the emerging hypersynchronous connectivity during the generalized seizure is independent of spatial constraints of baseline or preictal connectivity state. Furthermore, functional connectivity patterns during generalized seizures exhibit no or little negative correlations that would be driven by inhibitory connections. All these drastic alterations in brain activity and connectivity during seizure propagation is in line with the general view of epileptic seizures as state transitions, where brain networks shift from a balanced state to a hypersynchronous or hyperconnected state (Kramer and Cash, 2012; Kramer et al., 2008; Ponten et al., 2007; Schindler et al., 2008). Intriguingly, epileptic seizures in human patients are more likely to occur during transitions of sleep states which show profound alterations in brain synchrony and connectivity (Herman et al., 2001; Hofstra et al., 2009). Similar to epileptic state transitions, higher frequency desynchronized activity during wakefulness and REM sleep is replaced by low-frequency oscillations during slow wave sleep (Harris and Thiele, 2011). Moreover, thalamocortical and corticocortical pathways that are involved in sleep (Bazhenov et al., 2002; Steriade et al., 1993) can facilitate long range connectivity and are proposed to play important roles in epileptic seizures (Beenhakker and Huguenard, 2009; Paz et al., 2013; Paz and Huguenard, 2015). Hence, investigating parallels between sleep state transitions and epileptic state transitions are promising avenues for better understanding of common mechanisms that might be used by these processes. Given the drastic changes in neuronal connectivity during generalized seizures, we propose that interfering with this process might require strategies that involve manipulations beyond functional connectivity diagrams in healthy brains.

The rapid transition of neuronal activity and connectivity that we observed was difficult to explain by only relying on the rules of neuronal connectivity mediated by classical synaptic transmission. The rapid and excessive increase in positive correlations that is not constrained by spatial distances between neurons, encouraged us to investigate the properties of non-neuronal cells, such as glia, during seizure generation. Our results showed overwhelming differences between the activity patterns of glial and neuronal networks. Already during the preictal phase, glial networks were highly active and synchronized across large distances, when compared to rather small and local increase of neuronal activity and synchrony. Moreover, we observed very little functional interactions between glial and neuronal network activity during baseline and preictal state. Yet, this picture completely changed during generalized seizures, that were characterized by a rapid increase in functional coupling of glial and neuronal networks. These results suggest that the initiation of generalized seizures can be mediated by a transient functional coupling between glial and neuronal networks.

Astrocytes are known to regulate the availability of neurotransmitters and ions at the synaptic cleft, thereby modulating synaptic transmission and neuronal excitability (Bazargani and Attwell, 2016; Perea et al., 2009; Volterra et al., 2014). Moreover, the role of glia in the generation and prevention of epileptic seizures is an expanding field. Several mutations in glia associated genes were linked with epilepsy (Guella et al., 2017; Wallraff et al., 2006). Especially, gap junction coupling between astrocytes was shown to play important roles in distribution and homeostasis of ions and neurotransmitters across large distances in the brain (Allen and Eroglu, 2017; Alvarez-Ferradas et al., 2015; Bazargani and Attwell, 2016; Carmignoto and Haydon, 2012; Coulter and Steinhauser, 2015; Eid et al., 2007; Perea et al., 2009; Scemes and Spray, 2004; Schroder et al., 2000; Steinhauser et al., 2012; Volman et al., 2012; Volterra et al., 2014; Wetherington et al., 2008). In fact, our results revealed strong functional connections within glial networks that can span several hundred micrometers across the brain. These findings are in line with the idea that glial networks are strongly coupled and can act as a single unified entity with highly coordinated activation (Steinhauser et al., 2012). Surprisingly during preictal state, we observed that the highly synchronous and strong glial activity appears to be functionally uncoupled from neuronal activity for long periods of time. During this period, the neuronal activity is relatively balanced and does not present large changes, except regional synchronous network activity. We propose that during this preictal state, large synchronous glial activity represents the homeostatic function of glia that can absorb excessive levels of glutamate and cations through the major glial excitatory amino acid transporter, EAAT2 (also known as glutamate transporter 1, GLT1; and solute carrier family 1 member 2, SLC1A2) (Pines et al., 1992), and redistribute these molecules and cations across the gap junction coupled glial networks (Danbolt, 2001), balancing neuronal activity. In line with this hypothesis, *GLT1* knockout mice display spontaneous seizures (Tanaka et al., 1997) and mutations in *SLC1A2* gene are associated with epilepsy in human patients (Guella et al., 2017; Myers et al., 2016). Therefore, we propose that synchronized preictal glial activity in our recordings reflects the protective function of glia against epileptic seizures. Interestingly, it was reported that gap junction coupling between astrocytes is diminished during epileptogenesis and results in a reduced homeostatic potential of astrocytes (Bedner et al., 2015; Steinhauser et al., 2012; Strohschein et al., 2011; Wallraff et al., 2006). Moreover, deficiency in the astrocytic gap junction coupling was shown to cause spontaneous seizures (Wallraff et al., 2006) and temporal lobe epilepsy (Bedner et al., 2015; Schroder et al., 2000; Strohschein et al., 2011; Wallraff et al., 2006).

Given all these protective effects of glia against epileptic seizures, the role of strong coupling between glial and neuronal networks during seizure generation remains elusive. We propose that during prolonged periods of preictal activity the excessively used homeostatic function of glia might reach to saturation or fatigue. This leads to an eventual collapse of homeostatic glial function, thereby releasing the glial content especially the glutamate, delivering massive excitation across the brain and triggering a generalized seizure. The cellular mechanism for the collapse of homeostatic glial function and the precise trigger of the generalized seizures are yet to be identified. Another potential mechanism that could explain such rapid alterations in glial function might be directly related to the way the major glial glutamate transporter, EAAT2, operates (Danbolt et al., 1992; Kim et al., 2011; Tanaka et al., 1997). EAAT2 uptakes extracellular glutamate by exchanging it with intracellular potassium ions (Danbolt, 2001). Hence, the clearing of excessive glutamate in the preictal brain is a process heavily dependent on intra-and extracellular potassium levels. It is likely that during preictal activity extracellular potassium levels reach a threshold, which prevents EAAT2 from functioning. The glial network may eventually collapse, triggering a massive release of glutamate from the strongly coupled glial network, initiating a generalized seizure. In fact, our optogenetic activation of small number of glial cells in zebrafish brain was very effective in generating transient and strong activation of nearby neurons, highlighting a potential way to excite neurons through gliotransmission (Haydon, 2001; Parpura et al., 1994; Volterra and Meldolesi, 2005). Further experiments will be important for a better mechanistic understanding of the role of glial networks both in prevention and induction of epileptic seizures.

While in this study we focused on the function of glia-neuron interactions during the transitions from preictal to generalized seizure states, these interactions are likely to be involved in many other types of state transitions within the brain networks. An expanding number of studies propose diverse roles for glia-neuron interactions in brain functions such as regulating locomotion, memory formation, arousal and sleep (Adamsky et al., 2018; Brancaccio et al., 2017; Nimmerjahn et al., 2009; Slezak et al., 2018). Given the large number of glial cells in the brain, their prominent role in regulating neuronal excitability and activity is not that surprising. Future studies on the mechanistic understanding of glia-neuron communication and the plasticity of these connections will shed further light on common mechanisms between the physiology and pathophysiology of brain circuits.

## ACKNOWLEDGEMENTS

We thank K. Kawakami (NIG-Mishima, Japan), M. Ahrens (HHMI, Janelia Farm, USA), C. Wyart (ICM, Paris, France), Michael Orger (Champalimaud Centre for the Unknown, Lisbon, Portugal) and H. Baier (MPI, Martinsried, Germany) for transgenic lines, T. Ohshima (Waseda university, Tokyo, Japan) for the *GFAP:Gal4* plasmid. We thank M. Andresen, V. Nguyen and our fish facility support team for technical assistance. We also thank E. Brodtkorb (St Olav’s University Hospital and NTNU) and the Yaksi lab members for stimulating discussions.

This work was funded by The Liaison Committee for Education, Research and Innovation in Central Norway (‘Samarbeidsorganet’) Grant (S.M.-S., N.J.-Y., E.Y.), Flanders Science Foundation (FWO) Grant (C.D.V., E.Y.), RCN FRIPRO Research Grant 239973 (E.Y.) and ERC starting grant 335561 (R.P., E.Y.). Work in the E.Y. lab is funded by the Kavli Institute for Systems Neuroscience at NTNU.

## AUTHOR CONTRIBUTIONS

*Conceptualization, E.Y., N.J.-Y.; Methodology, C.D.V., S.M.-S., N.J.-Y., E.Y..; Data Analysis, C.D.V., S.M.-S., C.D., E.Y.; Software, C.D.V., R.P.; Providing reagents A.M., K.K.; Investigation, all authors; Writing, C.D.V., S.M.-S., N.J-Y., E.Y.; Review & Editing, all authors; Funding Acquisition, S.M.-S., N.J.-Y., E.Y.; Supervision, N.J.-Y., E.Y*.

## STAR METHODS

### Contact for Reagent and Resource Sharing

Further information and requests for reagents may be directed to, and will be fulfilled by the lead author Emre Yaksi.

### Experimental Model and Subject Details

#### Fish maintenance

Fish were kept in 3.5 liter tanks at a density of 15-20 fish per tank in a Techniplast Zebtech Multilinking system at constant conditions: 28 °C, pH 7, 6.0 ppm O_2_ and 700 µS, at a 14:10 hour light/dark cycle to simulate optimal natural breeding conditions. Fish received a normal diet of dry food (Zebrafeed, Sparos I&D Nutrition in Aquaculture, <100-600, according to their size) two times per day and *Artemia nauplii* once a day (Grade0, platinum Label, Argent Laboratories, Redmond, USA). Larvae were maintained in egg water (1.2 g marine salt, 20 L RO water, 1:1000 0.1% methylene blue) from fertilization to 3 dpf and in artificial fish water (AFW, 1.2 g marine salt in 20 L RO water) from 3 to 5 dpf. The animal facilities and maintenance of the zebrafish, *Danio rerio*, were approved by the Norwegian Food Safety Authority (NFSA) and Belgian government. All experimental procedures performed on zebrafish larvae up to 5 days post fertilization were in accordance with the directive 2010/63/EU of the European Parliament and the Council of the European Union and the Norwegian Food Safety Authorities. Experimental procedures performed on zebrafish larvae older than 5 dpf were further approved by the Ethical Committee of KULeuven in Belgium and Norwegian Food Safety Authority.

For experiments, the following fish lines were used: *Tg(elavl3:GCaMP6s)* (Vladimirov et al., 2014), *Tg(elavl3:jRCaMP1a)* (Dunn et al., 2016), *Tg(UAS:ChR2-mCherry)* (Hubbard et al., 2016; Schoonheim et al., 2010), *Tg(gad1:GFP)* (Satou et al., 2013), *Tg(vglut2a:dsRED*) (Miyasaka et al., 2009), *Et(−0.6hsp70l:Gal4-VP16)s1020t* (Scott and Baier, 2009), *Tg(UAS:GCaMP6s*) (Muto et al., 2017). *Tg(GFAP:Gal4)nw7Tg* transgenic animals were generated in our lab upon coinjection of tol2 transposase mRNA (Kawakami et al., 2004; Reiten et al., 2017) and *GFAP:Gal4* plasmid obtained from the Ohshima lab (Shimizu et al., 2015). The *Tg(GFAP:Gal4)nw7Tg* expression pattern was identified in three independent founders.

#### Combined epifluorescence calcium imaging and LFP experiments

Simultaneous epifluorescence calcium imaging and LFP experiments were performed on 7 days old *Tg(elavl3:GCaMP6s)* zebrafish (n=5). Zebrafish larvae were first anesthetized with 0.02% MS222 and paralyzed by α-bungarotoxin injection (Dreosti et al., 2014; Vendrell-Llopis and Yaksi, 2015). They were embedded in 1% low melting point agarose (LMP, Fisher Scientific) in a recording chamber (Fluorodish, World Precision Instruments). The layer of LMP agarose covering the dorsal side of the forebrain was carefully removed, so it was accessible for electrode placement. The fish were placed under the epifluorescence microscope (Olympus BX51 fluorescence microscope, Olympus Corporation). AFW was constantly perfused during the experiments. For electrophysiological recordings, a borosilicate glass patch clamp pipette (9-15 MOhms) loaded with teleost artificial cerebrospinal fluid (Mathieson and Maler, 1988) (ACSF, containing in mM: 123.9 NaCl, 22 D-glucose, 2 KCl, 1.6 MgSO_4 ×_ 7H_2_O, 1.3 KH_2_PO_4_, 24 NaHCO_3_, 2 CaCl_2 ×_ 2H_2_O) was inserted into the optic tectum. Electrical recordings were performed in current clamp mode with a high impedance amplifier and band pass filtered at 0.1–1000 Hz, at a sampling rate of 10 KHz (MultiClamp 700B amplifier, Axon instruments, USA). Calcium signals were recorded using an EMCCD camera (Hamamatsu Photonics) at sampling rate of 25 Hz. After electrode placement the larvae were left 10 minutes for stabilization to ensure fish were fully awake and recovered from the anesthesia, MS222. Baseline activity was recoded for 10 minutes, then 20 mM pentylenetetrazole (PTZ) dissolved in AFW was delivered via perfusion system until the end of the experiment. Duration of the total recording was 90 minutes. Data acquisition of both signals (LFP and calcium) were performed with a custom code written in MATLAB (Mathworks).

#### Confocal anatomical imaging

For confocal anatomical imaging, 5 days old *Tg(GFAP:Gal4)nw7Tg;Tg(elavl3:jRCaMP1a) and Tg(gad1:GFP);Tg(vglut2a:dsRED*) zebrafish larvae were anesthetized with 0.02% MS222 and embedded in 1.5-2% LMP agarose. Anatomical Z-scans were acquired using a Zeiss Examiner Z1 confocal microscope with a 20x plan NA 0.8 objective, using 10x average for each plane. Counting of GABAergic and glutamatergic neurons was done semi-automatically with a custom code written in MATLAB (Mathworks).

#### Combined optogenetical stimulation and electrophysiological recordings

For combined optogenetical stimulation and electrophysiological recordings, 4-5 weeks old zebrafish were anesthetized in ice cold water and sacrificed by decapitation. Their brains were dissected out in cooled and oxygenated ACSF. For intracellular recordings, borosilicate glass capillaries of 9-15 MOhms were filled with intracellular solution which contained (in mM): 130 K-gluconate, 10 Na-gluconate, 10 HEPES, 10 Na^2+^-Phospho-Creatine, 4 NaCl, 4 ATP-Mg and 0.3 Na^3+^-GTP. Electrical signals were recorded by MultiClamp 700B amplifier at sampling rate of 10 KHz. All recordings and data analyses were performed using custom codes written in MATLAB. Activation of channelrhodopsin2 was done by flashing a 480 nm LED light, through the optical path of the Olympus BX51 microscope for a duration of 1 second. For morphological reconstructions, neurobiotin (0.5%, Vectorlabs) was added to the intracellular solution. Brains were fixed overnight in 4% PFA at 4 °C, and afterwards incubated first with PBS and later with 5% streptavidin (Vectorlabs) in 0,5% PBSTx. Brains were imaged using a Zeiss Examiner Z1 confocal microscope.

#### Two-photon calcium imaging

Two-photon calcium imaging was performed on 7 days old *Tg(elavl3:GCaMP6s)* (n=8), 5 days old *Et(−0.6hsp70l:Gal4-VP16)s1020t);Tg(UAS:GcaMP6s)* (n=4) and 5 days old *Tg(GFAP:Gal4) nw7Tg;Tg(UAS:GCaMP6s)* zebrafish larvae (n=11). Animals were paralyzed with α-bungarotoxin (Invitrogen BI601, 1 mg/mL) and embedded in 1.5-2% LMP agarose in a recording chamber (Fluorodish, World Precision Instruments). The recordings were performed in two-photon microscopes (from Thorlabs Inc and Scientifica Inc) using a 16x water immersion objective (Nikon, NA 0.8, LWD 3.0, plan). A mode-locked Ti:Sapphire laser (MaiTai Spectra-Physics) tuned to 920 nm was used for excitation. Either single plane or volumetric recording (6 planes with a Piezo) were obtained. Acquisition rates were for the recordings of *Tg(elavl3:GCaMP6s)*, and *Et(−0.6hsp70l:Gal4-VP16)s1020t);Tg(UAS:GCaMP6s)* fish: 40 Hz for a single plane of 1408×384 pixels or 3 Hz for a volume of 1536×900 pixels × 6 planes. For *Tg(GFAP:Gal4)nw7Tg;Tg(UAS:GCaMP6s)* fish the acquisition rate was 24 Hz for a single plane of 1536×650 pixels. After a 3 minute baseline period, a solution of 20 mM PTZ diluted in AFW was added through a perfusion system in order to induce epileptic activity. Total duration of the recordings were 60-70 minutes. Data analysis was done with a custom code written in MATLAB (Mathworks) as described in the following section.

#### Data analysis

Two-photon microscopy images were aligned using an adapted algorithm (Reiten et al., 2017) that correct for occasional drift in the XY dimension, based on “hierarchical model-based motion estimation (Bergen et al., 1992). Individual neurons were automatically detected using a pattern recognition algorithm adapted from Ohki et al (Ohki et al., 2005), which identify neurons by using a correlation based approach comparing the *GCaMP6s* labeled neurons with torus or ring shaped neuronal templates (Jetti, 2014). Once all neurons were segmented they were individually tracked over the length of the whole recording to assure that the same neuron was properly captured. In case a neuron was lost during the tracking because of Z-drift, it was discarded from further analysis. Later, all automatically detected neurons are confirmed by visual inspection. Once the pixels belonging to each neuron were identified, the average of those pixels per image was calculated providing the complete time course of each individual neuron over time. Sparsely labelled glial cells were manually identified for Figure 5 and large numbers of glial cells analyzed in Figure 4 were semi-automatically detected as described above for neurons. For each cell, the fractional change in fluorescence (ΔF/F) relative to the baseline was calculated(Jetti, 2014). All traces from neurons and glia were resampled to a final rate of 4 Hz (using decimate function in MATLAB). Neuronal and glial activity was studied in windows of 3 minutes: baseline (3 minutes prior to drug delivery), preictal period (3 minutes preceding the seizure) and seizure. In each interval a cell was considered active if its change in fluorescence was greater than 3 times the standard deviation from the baseline period. Generalized seizure onset was manually annotated as the time point with sudden increase of neuronal activity recruiting the majority of the cells in all brain regions. The activity of each cell was calculated using the integral under the curve using trapezoidal numerical integration method (function trapz, MATLAB).

### Quantification and Statistical Analysis

Statistical analysis was done using MATLAB. Wilcoxon ranksum test was used for non-paired analysis and Wilcoxon signed rank test for paired analysis. P<0.05 was considered as statistically significant.

### Data and Software Availability

All data and software are available up on request

